# Axillary gland gene expression and potency reveal the nature of mimetic relationships in Corydoradinae catfishes

**DOI:** 10.64898/2026.01.27.701968

**Authors:** Emily Phelps, Ellen Bell, Simone Immler, Martin I. Taylor

## Abstract

The evolution of Müllerian or Batesian mimicry depends on the relative unpalatability of participating taxa, yet the mechanistic basis of such unpalatability remains poorly explored in many systems. Unpalatability in the Corydoradinae catfishes, known for their striking mimetic diversity, is underpinned by sharp, lockable spines, armoured bodies and the production of venoms/toxins. Here, we assess the contribution of the axillary gland, a structure at the base of the pectoral spine, to toxicity in two Corydoradinae genera, *Corydoras* and *Hoplisoma*. Using a brine shrimp cytotoxicity assay, we demonstrate that axillary gland extracts are significantly more toxic than muscle extracts in both genera, but toxicity does not differ significantly between genera. Transcriptomic analyses identified 539 candidate toxin genes upregulated in axillary gland tissue, relative to scute tissue, containing signal peptides and had predicted toxin functions based on amino acid sequences. Notably, candidate genes include domains typical of piscine venoms, such as lectins and peptidase S1. Although significant differences in gene expression were detected between genera in candidate toxins, log-fold changes were small, and predicted toxin potency was not significantly different between genera. Together, our findings indicate that axillary gland toxins are likely to contribute to unpalatability in Corydoradinae catfishes and provide support for Müllerian mimicry in this system.

## Background

Protective mimicry occurs when a close phenotypic resemblance in aposematic colouration between sympatric species has evolved to educate or deceive predators (Caro, 2017; Ruxton *et al*., 2018). One or more of the mimetic species may be toxic, venomous or otherwise unprofitable to predators rendering them unpalatable, for example, through sharp spines or bad taste (Ruxton *et al*., 2018; Caro & Ruxton, 2019). The nature and degree of this unpalatability is a key but understudied component of mimicry (Ruxton *et al*., 2018) and distinguishes Batesian and Müllerian mimicry. In Batesian mimicry, a palatable species mimics a non-palatable species. In Müllerian mimicry, both species are unpalatable and utilise a shared resemblance to deter predators (Bates, 1862; Joron & Mallet, 1998; Caro, 2017; Caro & Ruxton, 2019). In Müller’s original theory, co-mimics were considered equally unpalatable (Müller, 1878; Ruxton *et al*., 2018). However, one co-mimic may be more toxic/unpalatable than the other and this theoretically changes the mutualistic relationship to a parasitic one (quasi-Batesian mimicry), as the less toxic species erodes the protection of the more toxic species (Speed, 1993; Aubier *et al*., 2017; Ruxton *et al*., 2018). This suggests protective mimicry may be a continuum between Batesian mimicry and Müllerian mimicry rather than a strict dichotomy. However, whilst quasi-Batesian mimicry has been shown to be theoretically possible, it has never been demonstrated in an empirical system (Speed, 1993; Rowland *et al*., 2010).

Mimetic interactions can evolve via two main routes: i) advergence where one taxon evolves to resemble another or ii) convergence where both taxa converge on a shared phenotype (Symula *et al*., 2001; Ruxton *et al*., 2018). While Batesian (and quasi-Batesian) mimicry evolves through advergence, the evolutionary steps underlying Müllerian mimicry are less clear (Ruxton *et al*., 2018) and most theoretical evidence suggests it is driven by advergence. However, initial advergence followed by convergence (Sherratt *et al*., 2009; Merrill *et al*., 2015; Ruxton *et al*., 2018) or convergence alone (Hoyal Cuthill *et al*., 2019) can also underpin Müllerian mimicry. As such, the nature of the mimetic interaction, or location on the mimicry continuum, could provide an insight to the evolution of a mimetic system.

Toxin production in fishes can render them unpalatable via several pathways, including the synthesis of toxins within tissues (poison) or secretion of toxins from specialized glands, which may lack (ichthyocrinotoxins) or possess specialized delivery systems (venom) (Liu *et al*., 2025). Unpalatability in fish occurs most often through venom, and is usually delivered through sharp spines (but see Caley and Schulter 2003; Casewell *et al*., 2013; Smith *et al*., 2016). Over 2,900 cartilaginous and ray-finned fish species are considered venomous, with venom having evolved independently 18 or 19 times (Smith & Wheeler, 2006; Harris & Jenner, 2019). Piscine venom is thought to have evolved via the co-option of ichthyocrinotoxins, which have immune as well as predator deterrence functions (Cameron & Endean, 1973; Gratzer *et al*., 2015; Harris & Jenner, 2019). Despite the number of toxin producing fish, piscine toxicity is underexplored, partially due to the lack of defined venom or ichthyocrinotoxin gland structure and the resulting difficulties in extracting toxins (Harris & Jenner, 2019; von Reumont *et al*., 2022). However, improvements in RNA sequencing reduces some of these logistical issues as small amounts of gland tissue can elucidate toxin composition and evolutionary origins for taxa with little prior information (Magalhães *et al*., 2006; de Oliveira Júnior *et al*., 2016; von Reumont *et al*., 2022).

Catfishes (Order Siluriformes) are a highly diverse group of teleost fishes, comprising over 3,600 species across more than 40 families. Most possess sharp dorsal and pectoral spines, many of which can deliver venom (Haddad & Martins, 2006; Wright, 2009, 2015). The evolutionary origin and composition of venom in catfishes is not fully understood, but secretory cells on the sheath of the spine are thought to be primarily responsible for venom production in some species, releasing proteinaceous secretions through puncture wounds (Reed, 1907; Whitear *et al*., 1991; Wright, 2009, 2015). Additionally, many species possess an axillary gland, located at the base of the pectoral spine, and this has also been proposed as a source of toxin production (Reed, 1907; Halstead *et al*., 1953; Birkhead, 1972; Cameron & Endean, 1973b). It remains unclear whether axillary gland secretions contribute directly to toxicity or serve other functions e.g. antimicrobial (Greven *et al*., 2006; Kiehl *et al*., 2006). This uncertainty in role is partly due to lack of a clear delivery pathway from axillary gland to spine (Wright 2015). However, secretions are frequently released under handling stress (Greven *et al*., 2006), which suggests a possible defensive role. Consequently, the ecological role and biochemical makeup of axillary gland secretions require further study (Whitear *et al*., 1991; Wright, 2015).

Here we explore whether the axillary gland contributes to unpalatability in the Corydoradinae catfish and the extent to which it shapes ecological interactions among co-mimics. The Corydoradinae are a speciose group of Neotropical armoured catfish found across South America, consisting of seven genera (Dias *et al*., 2024). Corydoradinae are defended by a thick armour along with lockable dorsal and venomous pectoral spines (Greven *et al*., 2006; Alexandrou *et al*., 2011). In the Corydoradinae subfamily, 24 Müllerian mimicry rings have been identified, with sympatric species almost always belonging to different genera (Alexandrou *et al*., 2011). Not all genera are equally likely to participate in mimicry rings, with species from *Corydoras, Brochis* and *Hoplisoma* being overrepresented in mimetic communities (Alexandrou & Taylor, 2011). Interestingly, a whole genome duplication event (WGD) is thought to have occurred at the base of *Hoplisoma* (Marburger *et al*., 2018). Gene duplications have been shown to increase toxin dosage, replenishment and functional diversity of toxin proteins in other taxonomic groups (Wong & Belov, 2012). As such, gene duplication could affect toxin potency and complexity in the Corydoradinae.

We used a complementary suite of methods to better understand the nature of axillary gland secretions and their potential toxicity by comparing members of *Hoplisoma* and *Corydoras*. We used a brine shrimp cytotoxicity assay to test whether the axillary gland secretion showed evidence of toxicity, indicating a role in unpalatability, and whether there were differences in toxicity between Corydoradinae genera. We used differential expression of gene models among tissues alongside *in silico* predictors to identify putative toxin genes in a single *Corydoras* species (*C. simulatus*) to better understand the biochemical composition of the axillary gland secretion. Finally, using a phylogenetically controlled cross-genus approach, we explored differences in the putative venom gene expression between three species of *Corydoras* and three species of *Hoplisoma*. These complimentary approaches assist in elucidating the relative potency of axillary gland secretions in the two genera and consequently, the nature of the mimetic interaction between Corydoradinae co-mimics.

## Materials and Methods

### Brine shrimp cytotoxicity assay

#### Venom extract preparation

Extracts of axillary gland and muscle tissue were prepared from two individual fish per extract with the same individuals used for both tissues. Fish were acquired through the aquarium trade in the UK. Muscle tissue was chosen as a control tissue for the potency assay due to the reduction of contamination from the epidermal mucus. Axillary gland and muscle tissue extracts were prepared following Wright (2009), using 8ml of physiological saline per 1g of tissue. This change was necessary to obtain the volume of extract needed for the assay. This was followed by an additional centrifuge step for 5 minutes at 4°C at 6000 rpm to create the final raw axillary gland extract. A total of 80 extracts were prepared using fish tissue (n=20 per tissue per genus), using a total of 80 individuals (n=20 per genus). Individual length was recorded to account for effect of size and was measured as distance from snout to the base of the caudal fin using manual callipers to the nearest 0.5mm. The average length of individuals used in each extract was subsequently calculated.

#### Brine Shrimp Assay

Brine shrimp nauplii were reared from cysts (ZM Fish Food and Equipment (Winchester, UK)) and hatched in artificial seawater, made using Instant Ocean Aquarium salt (ZM Fish Food and Equipment, Winchester, UK), at 26°C in 14/10 Light/Dark cycle. Subsequently, 10 (± 1-2) 24-hour old live brine shrimp were added to 100 wells across multiple 96-well plates (0.3 mL wells), empty wells and those on the edge of the plate were filled with saline solution to reduce edge effects. Excess saline was removed and replaced with 10μl of axillary gland extract in each well using a micro-pipette. Extracts and saline only controls were replicated over four plates with randomised placement, using Well Plate Maker (Borges *et al*., 2021). Each extract was used once due to the small volume produced. The plate was covered with clear microplate sealing film and incubated for 24hrs under constant light conditions at room temperature (20-22°C). After 24 hours, the number of dead nauplii were counted and mortality calculated as the number of dead nauplii divided by the total number of nauplii.

The effect of genus and tissue type on brine shrimp mortality was assessed using a glmer() from the lme4 (v.1.1.35.1)(Bates *et al*., 2015) package in R (v. 4.2.3)(R Core Team, 2025) with mortality as the dependent variable and tissue type and lineage as fixed effects. Individual identity, representing the source of each tissue extract, was included as a random effect.

Mortality was modelled as a binomial trait, with the total number of nauplii in each well used as a weighting to account for the proportional data. Model residuals showed no significant deviation or dispersion, as assessed using DHARMa (v. 0.4.6)(Hartig, 2022). Additional models including average axillary gland size and average body length as a fixed factor were tested but excluded due to poorer model fit based on Akaike Information Criterion (AIC). Pairwise comparisons were performed using the package emmeans (v. 1.10.3)(Lenth, 2024). To evaluate the effect of tissue type on mortality independently of genus an additional GLMM was run with mortality as a function of tissue type and extract ID as a random factor. Genus specific differences in gland size was investigated using a linear mixed effects model-lmer() (lme4, v.1.1.35.1(Bates *et al*., 2015) and lmerTest, v. 3.1.3*(Kuznetsova et al*., 2017)).

### Transcriptome Sample Preparation and Sequencing

#### Sample Acquisition, RNA extraction and Sequencing

Species included represent three independent mimicry rings, each with a *Corydoras* species (*C. narcissus*, n=2, *C. desana*, n= 3, *C. simulatus*, n=7) and a *Hoplisoma* species (*H. granti*, n= 4, *H. tukano*, n=2, *H. metae*, n=5). Individuals were euthanised and the axillary gland tissue excised and immediately flash frozen using liquid nitrogen. Additionally, scute tissue (the thick armoured skin of *the Corydoradinae*) was taken from *C. simulatus* to provide a between tissue comparison. RNA was extracted using an adapted RNAeasy micro or mini kit (Qiagen, Germany) protocol with a prior Trizol step as detailed (http://www.untergasser.de/lab/protocols/rna_prep_comb_trizol_v1_0.htm). Quality control and library preparation for Illumina sequencing were performed at Novogene Cambridge, using NovaSeq 6000 and each sample sequenced to 40 million reads.

#### Transcriptome assembly and Annotation

To reduce mapping quality biases, species-specific master transcriptomes were created, with each individual’s reads mapped to a transcriptome of its respective species. In *C. simulatus* the master transcriptome contained sequences from both the axillary gland and scute tissues. This was achieved by combining raw sequence files of each species, trimming reads, removing adapters and filtering using a phred score of 20 using cutadapt (v. 2.10)(Martin, 2011). The combined and filtered sequences were visually inspected using FastQC (v. 0.11.8)(Andrews, 2010), and *de novo* assembled into transcriptomes using Trinity (v. 2.11.0)(Grabherr *et al*., 2011) using default settings. ChimeraTE (v.1.2)(Oliveira *et al*., 2023) was used to remove chimeric transcripts, resulting from transposable element insertions, using species-specific repeat libraries (Bell *et al*., 2022). Transcriptome completeness was evaluated using BUSCO (v5.3.2; (2015)) against the Actinopterygii_odb10 database.

Finally, the assembled transcriptomes were annotated using trinnotate (v. 4.0.2)(Bryant *et al*., 2017) with Uniprot (UniProt Consortium, 2021) and Pfam (Mistry *et al*., 2021) databases using blastn (v. 2.10.1+)(Camacho *et al*., 2009) and hmmer (v.3.3)(http://hmmer.org/).

### Toxin gene identification

#### Differential Expression analysis

Individual raw reads were trimmed and filtered to remove adapters and reads with a phred score below 20, using cutadapt (v. 2.10)(Martin, 2011). Both the axillary gland and scute tissue were pseudo aligned to the *C. simulatus* transcriptome using Salmon (v.1.2.1) (Patro *et al*. 2017). Transcript abundance was quantified using Salmon (v.1.2.1)(Patro *et al*., 2017) using the align_and_estimate_abundance.pl script from Trinity, for each individual using the species-specific transcriptome. Transcript abundance was normalised by gene length using tximport (v. 1.26.1) (Soneson *et al*., 2015). Differential expression analysis was performed using DESeq2 (v. 1.38.3)(Love *et al*., 2014).

#### Selection Criteria for Candidate Genes

Multiple lines of evidence were used to assess which genes were strong toxin candidates. Firstly, we identified genes that were significantly upregulated in the axillary gland compared to the scute tissue. Secondly, SignalP (v. 6.0.)(Teufel *et al*., 2022) was used to identify genes containing signal peptides (SPs), suggesting they may be secretory proteins (Emanuelsson *et al*., 2007; Owji *et al*., 2018). All known peptides representing precursors to toxins contain signal peptides (Fry *et al*., 2009). To reduce computational load, only the transcripts with the highest expression for each gene were used. These transcripts were translated into amino acid sequences using Transdecoder (v. 5.5.0) (https://github.com/TransDecoder/TransDecoder), selecting the longest ORF for each transcript.

SignalP was then run on the translated sequences using the eukaryota dataset. In addition to SignalP, TargetP (v.2.0)(Emanuelsson *et al*., 2000; Almagro Armenteros *et al*., 2019) was used to infer the intended subcellular location of the candidate proteins. This utilised the same translated transcripts as SignalP, and the results were filtered to retain sequences with a signal peptide only, removing those intended for the mitochondria. Finally, ToxinPred2 (v. 2.0) (Sharma *et al*., 2022) was used to estimate the toxicity of our candidate genes using the same translated transcripts detailed above. We ran both the random forest amino-acid composition (RF-AAC) and hybrid models. The RF-AAC gives a score which is proportional to the toxicity of the protein, whilst the hybrid model uses homology and motif estimation as well as RF-AAC to predict whether the protein is toxic.

#### Hidden Markov Models

To identify if any of the candidate genes contained domains related to known animal toxins HMMs were built using a database of animal toxins obtained from Uniprot (Ahmad *et al*., 2025). This method has previously been used to identify venom in the yellow catfish (*Pelteobagrus fulvidraco*)(Xie *et al*., 2016a). Proteins were retrieved using the search terms “venom”, “toxin”, “venomous”, “poison”, “poisonous”, “noxious” and “toxic”. Only reviewed proteins from animalia were retained. These were grouped by protein family, with only families with greater than 10 representative sequences retained, resulting in 107 families. Per-family HMMs were created using hmmbuild in HMMER (v. 3.3) (hmmer.org). Transdecoder was used (as above) to extract the longest ORF of each *C. simulatus* transcript which were translated into amino acid sequences. The HMMs were used to identify transcripts with known venom domains in the *C. simulatus* transcriptome using hmmsearch from HMMER.

#### Gene network analysis

Weighted gene network analysis was performed to identify genes with correlated expression, to further identify toxin candidates as well as housekeeping genes (Haney *et al*., 2019). Comparative gene network expression on the axillary gland and scute tissue from *C. simulatus* was performed by first normalizing the read data using vst() from DESeq2 (v. 1.38.3)(Love *et al*., 2014). To estimate expression modules, the pickSoftThreshold() and blockwiseModules() functions from WGCNA (v.1.72.5)(Langfelder & Horvath, 2008) were utilised. Modules were filtered to retain those with a candidate toxin gene. To explore differential expression of modules between axillary gland and scute tissue, a linear model was fitted using lmFit() from limma (v.3.54.2)(Ritchie *et al*., 2015). To stabilise variability in the estimates, empirical Bayes moderation, via the eBayes() from limma was applied. To correct for multiple testing, the Benjamini-Hochberg (BH) method was applied using topTable() and calculated the mean expression across the significantly differentially expressed modules.

#### GO term enrichment

Gene Ontology (GO) term enrichment analysis was used to indicate the biological function of genes within the expression modules. Annotated gene models within the focal module were assigned gene names using idmapping on uniprot.org (Ahmad *et al*., 2025). GO term enrichment analysis was then performed using g:Profiler (web version)(Kolberg *et al*., 2023) against the *Danio rerio* database using default parameters.

### Among genera variation in toxicity

#### Differential Expression analysis

For between genus comparisons, we allocated genes from different Corydoradinae species to orthogroups. Transcripts were filtered by longest isoform using the get_longest_isoform_seq_per_trinity_gene.pl from Trinity to ensure genes were only represented once before running Orthofinder (v. 2.5.2)(Emms & Kelly, 2019). Annotations were assigned to orthogroups using the Swissprot blastp hit, or if missing, the top Swissprot blastx hit, allocated to the *C. simulatus* gene present in the group. When multiple hits were assigned, the lowest evalue and the highest percentage identity annotation was allocated. Gene level inferences were estimated using tx2gene() from tximport. Data were pre-filtered to remove orthogroups with low counts and were normalized via vst() from DESeq2.

To estimate differential expression of toxin candidates, while controlling for phylogenetic structure, the EVE model was used (. Shifts in expression of toxin orthogroups on each branch on the phylogenetic tree (resolved by Orthofinder) were tested using the twoThetaTest() from the evemodel library in R (Gillard *et al*., 2021). Corrections for multiple testing were applied using p.adjust() in base

R. Subsequently logfold change was estimated from the log2 of the mean normalized count of an orthogroup in *Corydoras* over the equivalent of *Hoplisoma*.

#### Predicted Protein Toxicity

To predict the toxicity of our candidate toxins among *Hoplisoma* and *Corydoras* species we selected all transcripts within our toxin orthogroups. We then filtered transcripts to retain those with the highest expression per gene within each species. These were then translated into amino acid sequences using transdecoder, retaining sequences with the longest open reading frame (ORFs), as detailed above. The toxicity of these predicted proteins was then estimated ToxinPred2, using the RF-AAC model, retaining only the sequences that passed the default threshold for toxicity.

To investigate whether there was a difference in predicted toxicity between our candidate toxins we then adopted a mixed modelling approach, using lmer(). Using toxicity as the dependent variable, we included genera and sequence length as fixed effects as well as orthogroup as a random effect.

## Results

### Potency assay

A significant effect of tissue type on the mortality of brine shrimp (estimate = 2.668, SE = 0.433, z = 6.160, p < 0.0001) was identified with axillary gland causing significantly higher mortality than muscle tissue (Figure 1). However, there was no significant effect of genus (estimate =-0.213, SE = 0.804, z =-0.265, p = 0.791) nor the interaction between genus and tissue (estimate = 0.477, SE = 0.646, z = 0.739, p = 0.460). Post hoc tests revealed a significant effect of tissue within both *Corydoras sp*. and *Hoplisoma sp*. but no effect of genus (Supplementary Table 2). The effect of tissue (independent of genus) was compared to saline controls. Similarly, axillary gland extracts were significantly more toxic to brine shrimp than muscle extracts (estimate =-2.878, SE = 0.327, *z*.ratio =-8.796, p < 0.0001). The mortality within the saline control was significantly lower than the axillary gland extract (estimate =-3.502, SE = 0.757, z.ratio =-4.623, p < 0.0001) but not significantly different from the muscle extract (estimate = -0.623, SE = 0.788, z.ratio =-0.791, p = 0.709). Gland size could impact the amount of secretion produced (de Roodt *et al*., 2016). The *Corydoras* individuals used in this study were significantly larger (43.9mm, SE = 0.576) than the *Hoplisoma* individuals (31.9mm, SE = 0.361, t = 17.612, df = 31.922, p < 0.0001). However, after controlling for body size, there was no significant effect of genus on axillary gland size (estimate=-0.002, SE=0.001, df=17.400, t=-1.673, p = 0.112).

**Figure 1.**
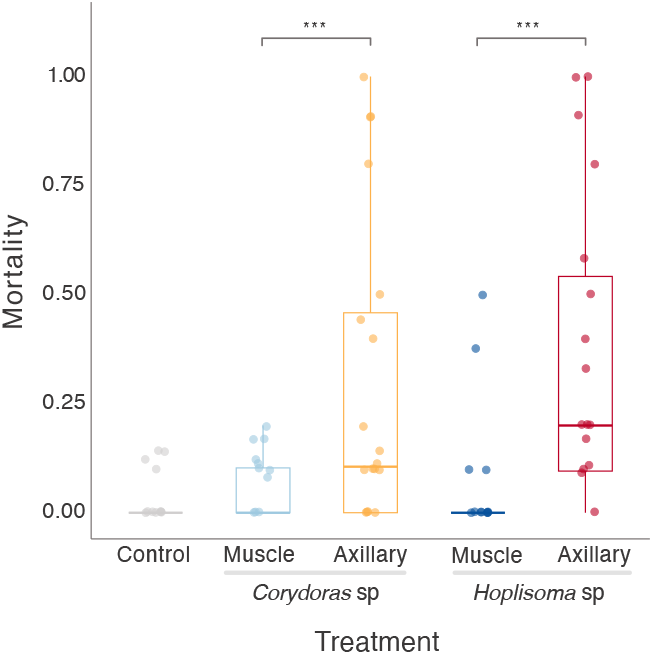
The effect of different tissue extracts and *Corydoradinae* genera on mortality on 24-hour old brine shrimp after a 24 hour incubation period. Asterisks denote significant differences, *** = p < 0.0001.

### Transcriptome Assembly

To identify candidate venom genes and quantify potential differences in expression between comimics, axillary gland (n = 6) and scute tissues (n = 5) of *C. simulatus* were sequenced. To enable a between genus comparison, the axillary gland of *H. granti* (n = 4), *H. metae* (n = 5), *H. tukano* (n = 2), *C. narcissus* (n = 2) and *C. desana* (n = 3) were also sequenced. The mean number of reads per individual across all species was 47,163,373 (range= 39,776,276-58,749,872), with similar mean read counts between scute (42,959,497) and axillary gland tissue (48,137,400), within *C* .*simulatus*. The *de novo* assembled transcriptomes showed good quality and had an average BUSCO score of complete gene models of 78.1%, indicating a good assembly and low fragmentation (Supplementary Table 1).

### Toxin gene identification

Within *C. simulatus*, samples clustered by tissue type (Supplementary Figure 1). A total of 5,812 genes were significantly differentially expressed (p_adj_ < 0.05) between the two tissues, representing 23% of genes. There were 2,746 genes upregulated in the axillary gland, relative to the scute tissue (Figure 2A). Of these genes, 539 met the criteria for being candidate toxin genes, including being estimated as toxic based on amino acid sequence, having a signal peptide and lacking a mitochondrial targeting sequence (Figure 2B, Supplementary File 1). Of these candidate genes, 29 contained domains similar to those from protein families known to be toxic (Figure 2C). This does not exclude candidates without these domains but can be informative to whether there is any convergence in the recruitment of proteins as toxins, as seen across animalia (Fry et al., 2009). The domains of the true venom lectins, snaclec and peptidase S1 families were most highly represented in the putative toxin genes (Figure 2C).

**Figure 2.**
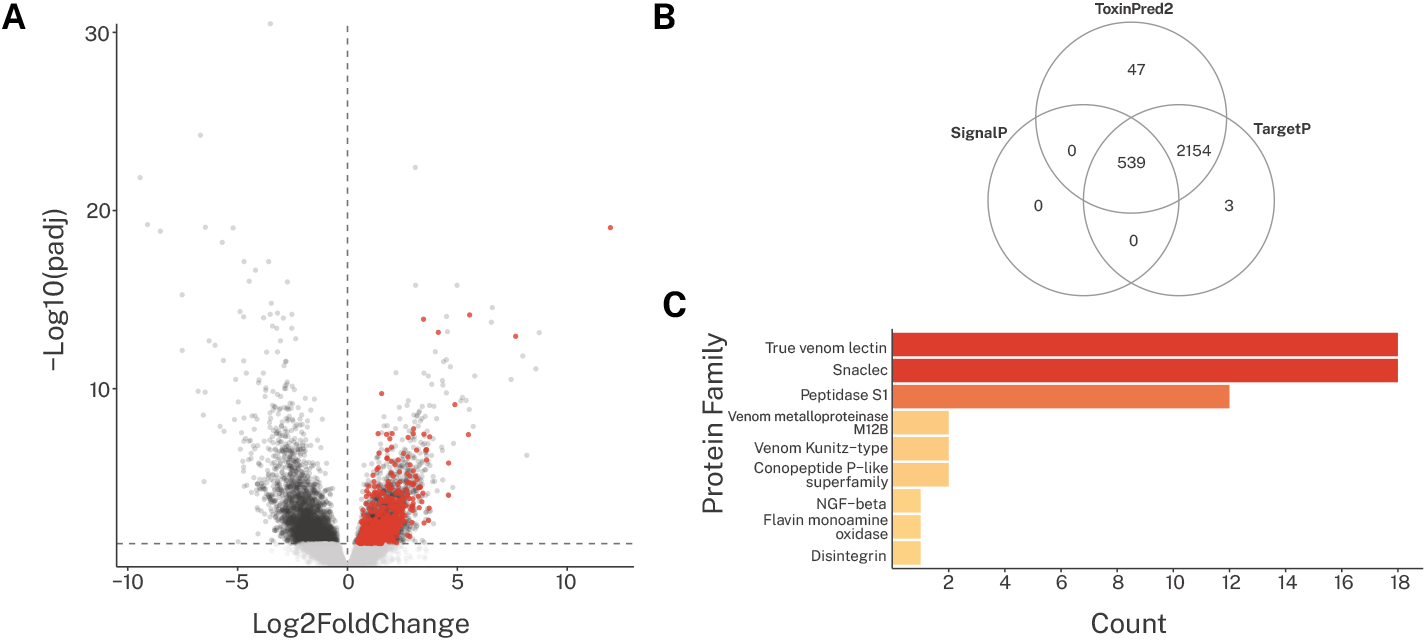
**A** Volcano plot showing the differential gene expression between *C. simulatus* axillary gland and scute tissue. The putative toxin genes are highlighted in red. **B** Selection criteria for candidate toxin genes, based on the estimated toxicity (ToxinPred2), presence of a signal peptide (SignalP) and the subcellular location of the protein (TargetP). **C** The frequency of toxin domains from known toxin protein families were identified in the candidate toxin genes using hidden Markov models (HMM).

Co-expression analysis identified eight modules containing candidate toxin genes that were significantly differentially expressed between the axillary gland and the scute tissue (Figure 3A-B, Supplementary Figure 2 and Supplementary Table 3). Three modules were significantly upregulated in the axillary gland relative to the scute tissue. These modules also contain the majority of toxin candidates (56.7%, Supplementary Table 2). Identified modules contained an additional 4,102 genes, which may be implicated in the generation and transportation of toxin proteins, representing putative toxin housekeeping genes (Supplementary File 1). Gene ontology enrichment showed modules contained genes relating to transmembrane transport (electron coupled proton transport, proton transmembrane transport and energy coupled proton transmembrane transport against electrochemical gradient), cellular respiration (ATP synthesis coupled electron transport, mitochondrial ATP synthesis, respiratory electron transport chain etc.), cellular organization and development (Cytoskeleton organization, anatomical structure morphogenesis, organelle organization etc.).

**Figure 3.**
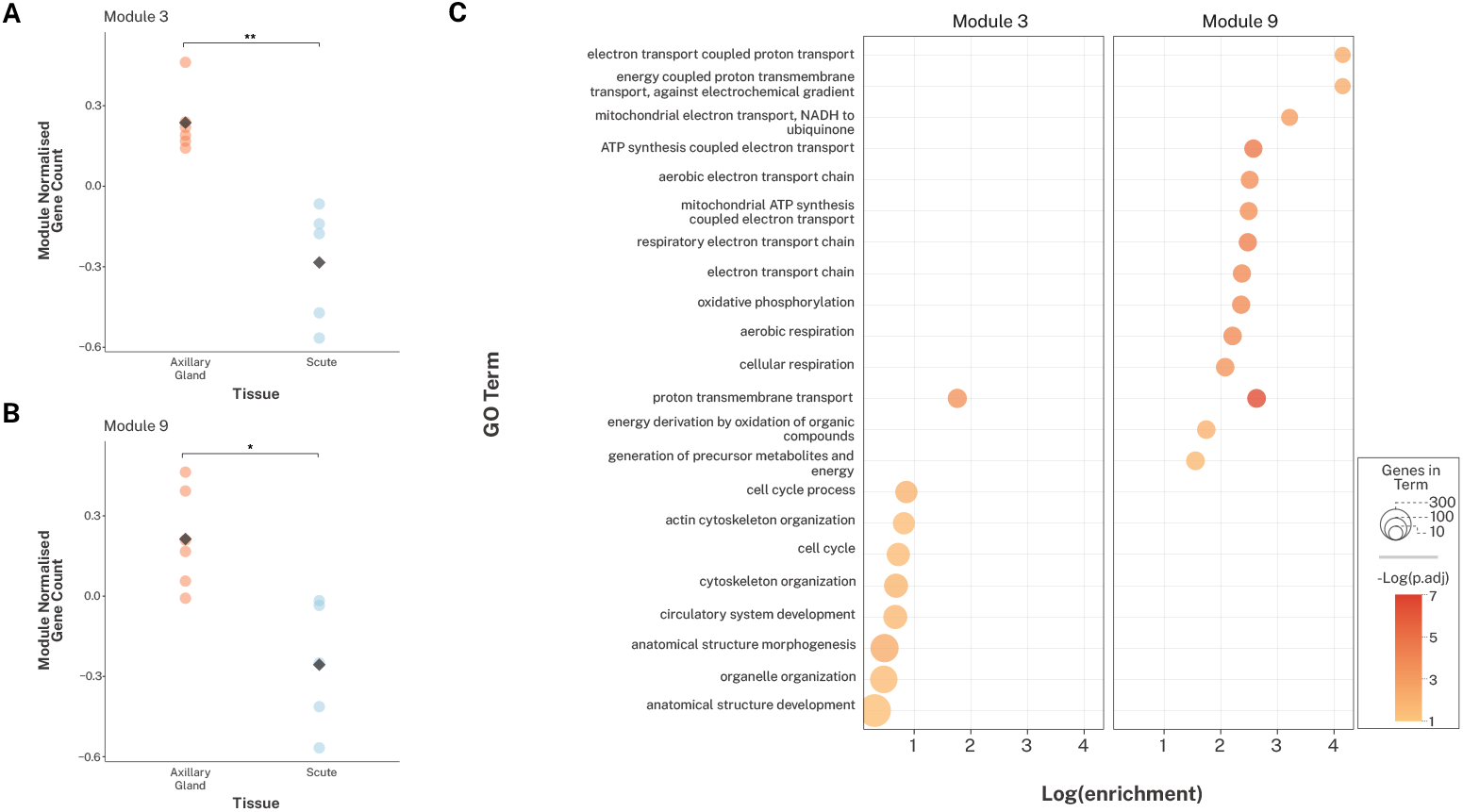
The toxin network modules represent a group of co-expressed genes thought to be involved in toxin production and regulation, including the majority of identified toxin genes. Difference in mean module expression between the axillary gland and scute tissue of *C. simulatus* in **A** Module 3 and **B** Module 9. Significant difference is denoted by asterisks (p <0.05, *, p <0.001 **). **B** Gene ontology (GO) enrichment of all genes in two of the toxin network modules. Module 8 was not included due to the lack of GO hits. Size indicates number of gene in the module with the associated GO term. Colour indicates statistical significance.

### Differential Expression in *Hoplisoma* and *Corydora*s axillary glands

We investigated the relative expression of toxin candidate orthogroups between *Corydoras* and *Hoplisoma*, which are often co-mimics. Samples clustered within species and genus (Supplementary Figure 3). Only 20 toxin candidate orthogroups exhibited a significant shift in expression (p < 0.05) at the branch delimiting *Corydoras* and *Hoplisoma* (Figure 4A). Whilst some of these genes showed an increase in expression in *Hoplisoma* relative to *Corydoras* (Figure 4A, C). However, of all the orthogroups identified, log2fold change was low (Supplementary Table 3).

**Figure 4.**
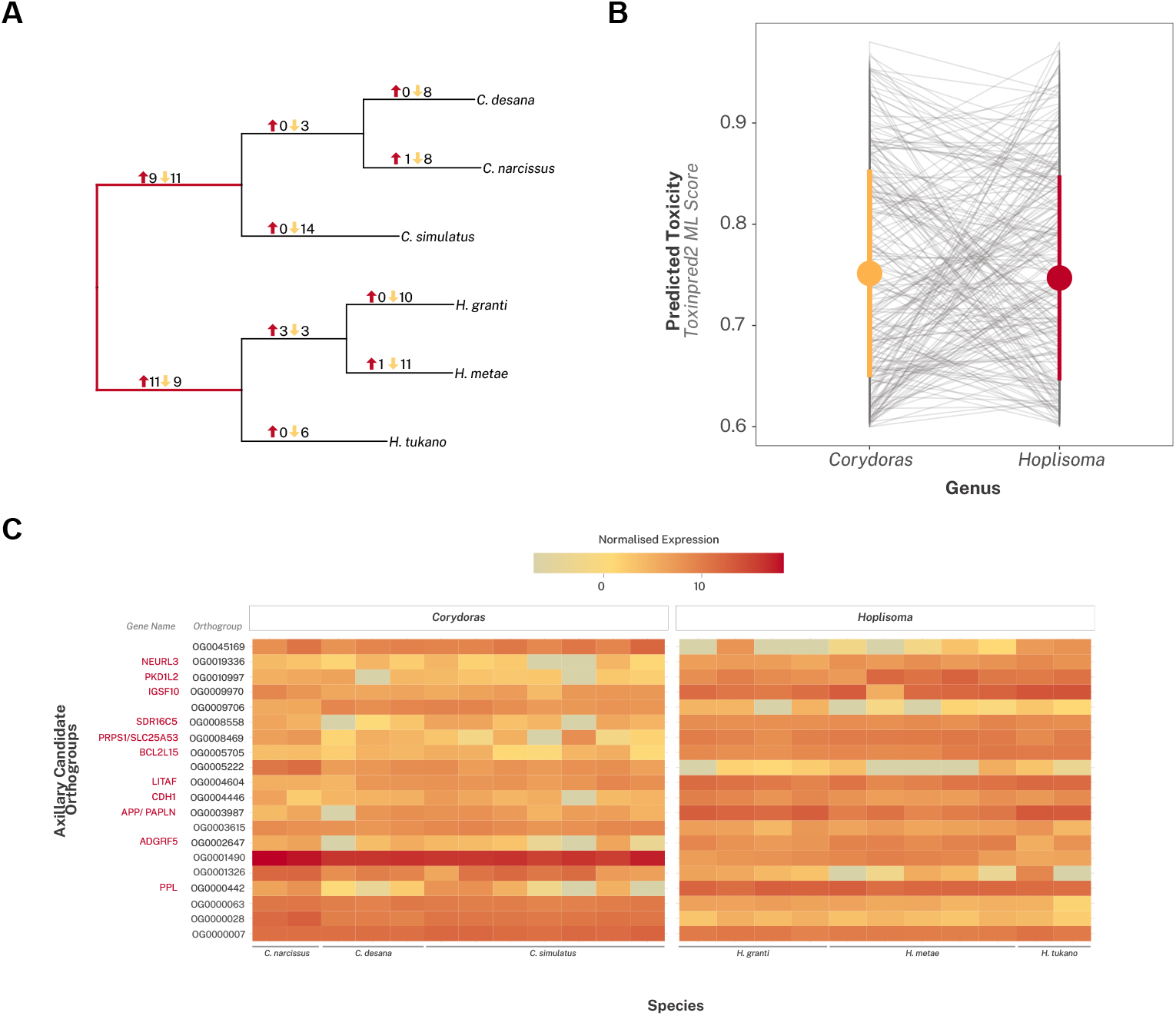
**A** Phylogeny of species used in this study, representing *Hoplisoma* and *Corydoras*. Numbers on Branches represent the number of genes with a shift in expression on that branch. Highlighted branch in red indicates the split between *Corydoras* and *Hoplisoma*. **B** Differences in the toxicity of candidates among genera, predicted by ToxinPred2. Coloured point represents the mean toxicity per genera whilst the coloured line represents the standard deviation. The grey lines link genes from the same orthogroup among the genera. **C** Heatmap of toxin orthologs with a shift in expression across the phylogeny in *Corydoras* species and *Hoplisoma* species. Orthogroups upregulated in *Hoplisoma* have been annotated to provide a gene name highlighted in red, corresponding to the highlighted red branches of the phylogeny. Each column represents a single individual. Expression data has been normalized, and species are given at the base of the heatmap.

### *In silico* toxin prediction in *Hoplisoma* and *Corydora*s

To explore whether *Hoplisoma* could be more toxic relative to *Corydoras* through increased toxin potency, we used RF-AAC to predict the potency of candidate toxins between the genera. We found no significant difference in the predicted potency of toxin candidates, indicating no difference between the groups (Figure 4B, p = 0.542). We included sequence length as a fixed effect to account for differences in transcript length among assembled transcriptomes but found no significant effect of length (p = 0.233). Our intercept was significant (p > 0.001), indicating a deviation from zero. This is perhaps due to only toxic genes (predicted toxicity > 0.6) being included in the model.

## Discussion

We explored the role of the axillary gland in Corydoradinae unpalatability to better understand protective mimicry in this system. We found axillary gland tissue extracts were significantly more toxic to brine shrimp than muscle tissue extracts, indicating that they may play an important role in predator deterrence. We then used RNA sequence data to identify candidate toxin genes alongside genes related to toxin production and maintenance within a species of *Corydoras, C. simulatus*. The presence of genes with characteristics of known animal toxins further indicated that the axillary gland plays a role in unpalatability within the group. Our brine shrimp lethality assay suggested there were no significant differences in toxicity of axillary gland extracts from *Corydoras* and *Hoplisoma*, two Corydoradinae genera which commonly form mimetic communities in South American rivers. This was supported by few candidate toxin genes having a significant shift in expression among the two genera. The candidate toxin genes which did demonstrate a shift in expression had a low logfold change indicating limited difference in expression. Overall, our results support the hypothesis that the axillary gland contributes to unpalatability and predator deterrence in the Corydoradinae, and that *Hoplisoma* and *Corydoras* represent Müllerian mimics, rather than quasi-Batesian mimics, with respect to the axillary gland.

Both the brine shrimp assay and RNA evidence support the use of the axillary gland in predator deterrence in the Corydoradinae. This is consistent with prior work, which showed that gland secretions resulted in self-poisoning (Kiehl *et al*., 2006). We explicitly tested for toxicity in the secretions using a brine shrimp lethality assay. Whilst brine shrimp are not the target species for Corydoradinae toxins, they give a strong indication that gland extracts are toxic. Corydoradinae are likely predated by both fish and avian predators (Alexandrou *et al*., 2011; Fuller, 2012). Brine shrimp assays have been used to test for toxicity of snake venoms as a proxy for mouse models, and there is evidence to show they do reflect toxicity to fish (Winters *et al*., 2018, 2022; Okumu *et al*., 2020, 2021; Chan *et al*., 2021). Whilst the gold standard to test for ecological relevance of toxicity in this gland is injection into/ingestion by a known predator, this is was not performed here for logistical and ethical reasons.

Corydoradinae fit the current descriptions of ichthyocrinotoxic fishes, which lack scales and use mucus as protection (Greven *et al*., 2006; Fuller, 2012; Dias *et al*., 2024; Liu *et al*., 2025). Ichthyocrinotoxins have been suggested to play a role in predator defence (Abdul-Haqq & Shier, 1991; Harris & Jenner, 2019; Lennox-Bulow *et al*., 2023). Ichtyocrinotoxic fishes are distinct from acanthotoxic (venomous fishes) through the lack of a delivery structure such as a spine (Liu *et al*., 2025). Whilst the dorsal spine of the Corydoradinae is considered a delivery apparatus, the pectoral spines are not, which has led researchers to classify the axillary gland as distinct from a specific venom gland (Wright, 2015). The principal reasons include the lack of a channel or route from the axillary gland to spine, which is not hollow, to transfer venoms and that the axillary gland secretion is water-soluble (Wright, 2015). Our evidence suggests the axillary gland secretions could contribute to unpalatability, even in the absence of a direct gland-to-spine delivery mechanism. Spines inflict physical damage on ingestion by predators. The axillary secretions, released by the Corydoradinae under stress, could enter these wounds, causing additional irritation and pain, subsequently encouraging prey release. As such, the combination of the spine and axillary gland secretion provides a plausible mechanism for predator deterrence, potentially acting in conjunction with the dorsal spine-based venom delivery.

To our knowledge, this is the first study to directly investigate the genes expressed in the axillary gland of a catfish. Some of the candidate toxin genes identified were found to have true venom lectins (C-type lectin), snaclec and peptidase S1 domains. Both snaclec domains and true venom lectin are C-type lectins, so both being present is unsurprising (Clemetson *et al*., 2009). C-type lectins have been identified as components of venom in the Scorpaeniformes (Scorpion fish)(Ellisdon *et al*., 2015; Campos *et al*., 2016; Ziegman *et al*., 2019). Animal toxins appear to repeatedly evolve from the same gene families, despite considerable phylogenetic distances between toxic taxa (Casewell *et al*., 2013). For example, the phospholipase A_2_ proteins are part of the venom arsenal of squamate reptiles, cephalopods and insects (Fry *et al*., 2009). Thus, the conserved domains of known toxin proteins offer a powerful framework for the identification of novel toxin genes (Fry *et al*., 2009; Xie *et al*., 2016b). Additionally, we have shown that housekeeping genes are enriched for protein transport and activity (Figure 3B). This is similar to the findings in the common house spider (*Parasteatoda tepidariorum*), where venom module genes were enriched for GO terms relating to intracellular signalling, transport and signal transduction pathways (Zhu *et al*., 2023).

Whilst the evidence presented in the current study suggests the Corydoradinae co-mimics are likely to be true Müllerian mimics with regards to their axillary gland secretions, other features of toxin production and delivery may still result in unequal unpalatability. For example, the orientation and degree of spine serrations, which show variation among Corydoradinae genera, could inflict more/less damage and associated pain (Dias *et al*., 2024). Additionally, the Corydoradinae possess dorsal venom glands accompanied by dorsal spines (Wright, 2009). Secretions from this gland, may differ from the axillary gland and may differ among genera. By further investigating these contributors to unpalatability, we can start to better understand the ecological interaction among co-mimics, and subsequently better understand how mimicry evolved in this group. *Corydoras* is the basal genera in the Corydoradinae subfamily, and toxicity of the axillary gland in this genera suggests the ancestral Corydoradinae produced toxins from the axillary gland (Dias *et al*., 2024). Additionally, axillary glands are present in non-Corydoradinae members of the Callichthyidae, further supporting this observation (Wright, 2009). The presence of toxicity in the wider family could indicate that Müllerian mimicry evolved because the cost of toxin production had already been overcome in the group (Ruxton *et al*., 2018; Harris & Jenner, 2019).

In conclusion, we provide evidence that axillary gland secretions in the Corydoradinae do have toxic properties and suggest that the gland may play an important role in predator deterrence and contribute to the evolution of mimicry in this system. This is supported by the identification of candidate toxin genes with known toxin domains, including true venom lectins and peptidase S1.

Limited expression and predicted protein toxicity differences between *Corydoras sp*. and *Hoplisoma sp*. are concordant with the potency assay, indicating that the axillary gland potency and complexity is conserved between the two genera. This work is an important contribution to the understanding of the genetics of mimicry, particularly of unpalatability, which is overlooked in favour of aposematic colouration.

## Data availability statement

Sequence data will be available on NCBI on acceptance of this manuscript; Brine shrimp toxicity data are available on Zenodo and code used for all analyses and plots available on github (https://github.com/EmilyPhelps/CorydoradinaeVenom2025).

## Ethics statement

Catfish samples were humanely euthanized under schedule 1 to the UK Animals (Scientific Procedures) Act 1986.

## Acknowledgements

We thank Ruth Sullivan for providing brine shrimp nauplii and Jayme Cohen for assistance with dissections. We would also like to thank Professor Tracey Chapman and Dr Domino Joyce for their insightful comments on early iterations of this manuscript.

## Author contributions

E.P: Conceptualized, Data curation, Formal analysis, Funding acquisition, Investigation, Methodology, Project administration, Resources, Visualisation, Writing-original draft, Writing-review and editing. M.I.T: Conceptualization, Funding acquisition, Supervision, Writing-review and editing. E.B: Supervision, Writing-review and editing. S.I: Supervision, Writing-review and editing.

## References

Abdul-Haqq, A.J. & Shier, W.T. 1991. Icthyocrinotoxins and Their Potential Use as Shark Repellents. J. Toxicol. Toxin Rev., doi: 10.3109/15569549109053859.

Ahmad, S., Jose da Costa Gonzales, L., Bowler-Barnett, E.H., Rice, D.L., Kim, M., Wijerathne, S., et al. 2025. The UniProt website API: facilitating programmatic access to protein knowledge. Nucleic Acids Res 53: W547–W553.

Alexandrou, M. & Taylor, M. 2011. Evolution, ecology and taxonomy of the Corydoradinae revisited. In: Identifying Corydoradinae CaJish: Aspidoras-Brochis-Corydoras-Scleromystax-C-numbers & CWnumbers, pp. 101–114.

Alexandrou, M.A., Oliveira, C., Maillard, M., McGill, R.A., Newton, J., Creer, S., et al. 2011. Competition and phylogeny determine community structure in Mullerian co-mimics. Nature 469: 84–88.

Almagro Armenteros, J.J., Salvatore, M., Emanuelsson, O., Winther, O., von Heijne, G., Elofsson, A., et al. 2019. Detecting sequence signals in targeting peptides using deep learning. Life Sci Alliance 2: e201900429.

Andrews, S. 2010. FastQC: A Quality Control Tool for High Throughput Sequence.

Aubier, T.G., Joron, M. & Sherratt, T.N. 2017. Mimicry among unequally defended prey should be mutualistic when predators sample optimally. Am. Nat. 189: 267–282.

Bates, D., Mächler, M., Bolker, B. & Walker, S. 2015. Fitting Linear Mixed-Effects Models Using lme4.

Bates, H.W. 1862. Contributions to an insect fauna of the Amazon valley. Lepidoptera: heliconinae. J. Proc. Linn. Soc. Lond. Zool. 6: 73–77.

Bell, E.A., Butler, C.L., Oliveira, C., Marburger, S., Yant, L. & Taylor, M.I. 2022. Transposable element annotation in non-model species: The benefits of species-specific repeat libraries using semi-automated EDTA and DeepTE de novo pipelines. Mol. Ecol. Res. 22: 823–833.

Birkhead, W.S. 1972. Toxicity of Stings of Ariid and Ictalurid Catfishes. Copeia 1972: 790–807.

Borges, H., Hesse, A.-M., Kraut, A., Couté, Y., Brun, V. & Burger, T. 2021. Well Plate Maker: a user-friendly randomized block design application to limit batch effects in large-scale biomedical studies. Bioinformatics 37: 2770–2771.

Bryant, D.M., Johnson, K., DiTommaso, T., Tickle, T., Couger, M.B., Payzin-Dogru, D., et al. 2017. A Tissue-Mapped Axolotl De Novo Transcriptome Enables Identification of Limb Regeneration Factors. Cell Rep. 18: 762–776.

Caley, M.J. & Schluter, D. 2003. Predators favour mimicry in a tropical reef fish. Proc. Biol. Sci. 270: 667–672.

Camacho, C., Coulouris, G., Avagyan, V., Ma, N., Papadopoulos, J., Bealer, K., 2009. BLAST+: architecture and applications. BMC Bioinformatics 10.

Cameron, A.M. & Endean, R. 1973a. Epidermal secretions and the evolution of venom glands in fishes. Toxicon 11: 401–410.

Campos, F.V., Menezes, T.N., Malacarne, P.F., Costa, F.L.S., Naumann, G.B., Gomes, H.L., et al. 2016. A review on the Scorpaena plumieri fish venom and its bioactive compounds. J. Venom. Anim. Toxins Incl. Trop. Dis. 22.

Caro, T. 2017. Wallace on coloration: Contemporary perspective and unresolved insights. Trends Ecol. Evol. 32: 23–30.

Caro, T. & Ruxton, G. 2019. Aposematism: Unpacking the defences. Trends Ecol. Evol. 34: 595–604.

Casewell, N.R., Wüster, W., Vonk, F.J., Harrison, R.A. & Fry, B.G. 2013. Complex cocktails: the evolutionary novelty of venoms. Trends Ecol. Evol. 28: 219–229.

Chan, W., Shaughnessy, A.E.P., van den Berg, C.P., Garson, M.J. & Cheney, K.L. 2021. The Validity of Brine Shrimp (Artemia Sp.) Toxicity Assays to Assess the Ecological Function of Marine Natural Products. J Chem Ecol 47: 834–846.

Clemetson, K.J., Morita, T. & Kini, R.M. 2009. Classification and nomenclature of snake venom C-type lectins and related proteins. Toxicon 54.

de Oliveira Júnior, N.G., Fernandes, G. da R., Cardoso, M.H., Costa, F.F., Cândido, E. de S., Garrone Neto, D., et al. 2016. Venom gland transcriptome analyses of two freshwater stingrays (Myliobatiformes: Potamotrygonidae) from Brazil. Sci. Rep. 6.

de Roodt, A.R., Boyer, L.V., Lanari, L.C., Irazu, L., Laskowicz, R.D., Sabattini, P.L., et al. 2016. Venom yield and its relationship with body size and fang separation of pit vipers from Argentina. Toxicon 121: 22–29. Elsevier BV.

Dias, A.C., Tencatt, L.F.C., Roxo, F.F., Silva, G. de S. da C., Santos, S.A., Britto, M.R., et al. 2024. Phylogenomic analyses in the complex Neotropical subfamily Corydoradinae (Siluriformes: Callichthyidae) with a new classification based on morphological and molecular data. Zool. J. Linn. Soc.

Ellisdon, A.M., Reboul, C.F., Panjikar, S., Huynh, K., Oellig, C.A., Winter, K.L., et al. 2015. Stonefish toxin defines an ancient branch of the perforin-like superfamily. Proc. Natl. Acad. Sci. U. S. A. 112: 15360– 15365.

Emanuelsson, O., Brunak, S., von Heijne, G. & Nielsen, H. 2007. Locating proteins in the cell using TargetP, SignalP and related tools. Nat Protoc 2: 953–971.

Emanuelsson, O., Nielsen, H., Brunak, S. & von Heijne, G. 2000. Predicting subcellular localization of proteins based on their N-terminal amino acid sequence. J Mol Biol 300: 1005–1016.

Emms, D.M. & Kelly, S. 2019. OrthoFinder: phylogenetic orthology inference for comparative genomics. Genome Biology 20: 238.

Fry, B.G., Roelants, K., Champagne, D.E., Scheib, H., Tyndall, J.D.A., King, G.F., et al. 2009. The toxicogenomic multiverse: convergent recruitment of proteins into animal venoms. Annu. Rev. Genomics Hum. Genet. 10: 483–511.

Fuller, I. 2012. Breeding Corydoradinae CaJish, 2nd ed. Ian Fuller Enterprises, Kidderminster, England.

Gillard, G.B., Grønvold, L., Røsæg, L.L., Holen, M.M., Monsen, Ø., Koop, B.F., et al. 2021. Comparative regulomics supports pervasive selection on gene dosage following whole genome duplication. Genome Biology 22: 103.

Grabherr, M.G., Haas, B.J., Yassour, M., Levin, J.Z., Thompson, D.A., Amit, I., et al. 2011. Full-length transcriptome assembly from RNA-Seq data without a reference genome. Nat. Biotechnol. 29: 644– 652.

Gratzer, B., Millesi, E., Walzl, M. & Herler, J. 2015. Skin toxins in coral-associated Gobiodon species (Teleostei: Gobiidae) affect predator preference and prey survival. Mar. Ecol. 36: 67–76.

Greven, H., Flasbecl, T. & Passia, D. 2006. Axillary glands in the armoured catfish Corydoras aeneus (Callichthyidae, Siluriformes). Verh. Ges. Ichthyol. 5: 65–69.

Haddad, V. & Martins, I.A. 2006. Frequency and gravity of human envenomations caused by marine catfish (suborder siluroidei): a clinical and epidemiological study. Toxicon 47: 838–843.

Halstead, B.W., Kuninobu, L.S. & Hebard, H.G. 1953. Catfish Stings and the Venom Apparatus of the Mexican Catfish, Galeichthys felis (Linnaeus). Trans. Am. Microsc. Soc. 72: 297–314.

Haney, R.A., Matte, T., Forsyth, F.S. & Garb, J.E. 2019. Alternative Transcription at Venom Genes and Its Role as a Complementary Mechanism for the Generation of Venom Complexity in the Common House Spider. Front Ecol Evol 7.

Harris, R.J. & Jenner, R.A. 2019. Evolutionary Ecology of Fish Venom: Adaptations and Consequences of Evolving a Venom System. Toxins 11.

Hartig, F. 2022. DHARMa: Residual Diagnostics for Hierarchical (Multi-Level / Mixed) Regression Models.

Hoyal Cuthill, J.F., Guttenberg, N., Ledger, S., Crowther, R. & Huertas, B. 2019. Deep learning on butterfly phenotypes tests evolution’s oldest mathematical model. Science Advances 5: eaaw4967.

Joron, M. & Mallet, J.L. 1998. Diversity in mimicry: paradox or paradigm? Trends Ecol. Evol. 13: 461– 466.

Kiehl, E., Rieger, C. & Greven, H. 2006. Axillary gland secretions contribute to the stress-induced discharge of a bactericidal substance in Corydoras sterbai (Callichthyidae, Siluriformes). Verh. Ges. Ichthyol. 5.

Kolberg, L., Raudvere, U., Kuzmin, I., Adler, P., Vilo, J. & Peterson, H. 2023. g:Profiler-interoperable web service for functional enrichment analysis and gene identifier mapping (2023 update). Nucleic Acids Res. 51: 207–212.

Kuznetsova, A., Brockhoff, P.B. & Christensen, R.H.B. 2017. lmerTest Package: Tests in Linear Mixed Effects Models.

Langfelder, P. & Horvath, S. 2008. WGCNA: an R package for weighted correlation network analysis. BMC Bioinformatics 9: 559.

Lennox-Bulow, D., Smout, M., Loukas, A. & Seymour, J. 2023. Stonefish (Synanceia spp.) Ichthyocrinotoxins: An ecological review and prospectus for future research and biodiscovery. Toxicon 236: 107329.

Lenth, R.V. 2024. emmeans: Estimated Marginal Means, aka Least-Squares Means.

Liu, L., Tang, X., Xu, J., Huang, Y., He, H. & Zhang, F. 2025. Ichthyotoxic fishes: a brief overview and prospectus for applications and future research. Toxin Reviews.

Love, M.I., Huber, W. & Anders, S. 2014. Moderated estimation of fold change and dispersion for RNA-seq data with DESeq2. Genome Biol. 15.

Magalhães, G.S., Junqueira-de-Azevedo, I.L.M., Lopes-Ferreira, M., Lorenzini, D.M., Ho, P.L. & Moura-da-Silva, A.M. 2006. Transcriptome analysis of expressed sequence tags from the venom glands of the fish Thalassophryne nattereri. Biochimie 88: 693–699.

Marburger, S., Alexandrou, M.A., Taggart, J.B., Creer, S., Carvalho, G., Oliveira, C., et al. 2018. Whole genome duplication and transposable element proliferation drive genome expansion in Corydoradinae catfishes. Proc. Roy. Soc. B 285.

Martin, M. 2011. Cutadapt removes adapter sequences from high-throughput sequencing reads. EMBnet.journal 17: 10–12.

Merrill, R.M., Dasmahapatra, K.K., Davey, J.W., Dell’Aglio, D.D., Hanly, J.J., Huber, B., et al. 2015. The diversification of Heliconius butterflies: what have we learned in 150 years? J. Evol. Biol. 28: 1417–1438. Wiley.

Mistry, J., Chuguransky, S., Williams, L., Qureshi, M., Salazar, G.A., Sonnhammer, E.L.L., et al. 2021. Pfam: The protein families database in 2021. Nucleic Acids Res. 49: 412–419.

Müller, F. 1878. Über die vortheile der mimicry bei schmetterlingen. Zoologischer Anzeiger 1: 4–55.

Okumu, M.O., Mbaria, J.M., Gikunju, J.K., Mbuthia, P.G., Madadi, V.O., Ochola, F.O., et al. 2021. Artemia salina as an animal model for the preliminary evaluation of snake venom-induced toxicity. Toxicon 12: 100082.

Okumu, M.O., Mbaria, J.M., Gikunju, J.K., Mbuthia, P.G., Madadi, V.O. & Ochola, F.O. 2020. Enzymatic activity and brine shrimp lethality of venom from the large brown spitting cobra (Naja ashei) and its neutralization by antivenom. BMC Res. Notes 13: 325.

Oliveira, D.S., Fablet, M., Larue, A., Vallier, A., Carareto, C.M.A., Rebollo, R., et al. 2023. ChimeraTE: a pipeline to detect chimeric transcripts derived from genes and transposable elements. Nucleic Acids Res 51: 9764–9784.

Owji, H., Nezafat, N., Negahdaripour, M., Hajiebrahimi, A. & Ghasemi, Y. 2018. A comprehensive review of signal peptides: Structure, roles, and applications. Eur. J. Cell Biol. 97: 422–441.

Patro, R., Duggal, G., Love, M.I., Irizarry, R.A. & Kingsford, C. 2017. Salmon provides fast and bias-aware quantification of transcript expression. Nat. Methods 14: 417–419.

R Core Team. 2025. R: A Language and Environment for Statistical Computing. Vienna.

Reed, H.D. 1907. The Poison Glands of Noturus and Schilbeodes. Am. Nat. 41: 553–566.

Ritchie, M.E., Phipson, B., Wu, D., Hu, Y., Law, C.W., Shi, W., et al. 2015. limma powers differential expression analyses for RNA-sequencing and microarray studies. Nucleic Acids Res. 43.

Rohlfs, R.V., Harrigan, P. & Nielsen, R. 2014. Modeling Gene Expression Evolution with an Extended Ornstein–Uhlenbeck Process Accounting for Within-Species Variation. Mol Biol Evol 31: 201–211.

Rowland, H.M., Mappes, J., Ruxton, G.D. & Speed, M.P. 2010. Mimicry between unequally defended prey can be parasitic: evidence for quasi-Batesian mimicry. Ecol. LeY. 13: 1494–1502.

Ruxton, G.D., Allen, W.L., Sherratt, T.N. & Speed, M.P. 2018. Avoiding AYack: The Evolutionary Ecology of Crypsis, Aposematism, and Mimicry. Oxford University Press.

Sharma, N., Naorem, L.D., Jain, S. & Raghava, G.P.S. 2022. ToxinPred2: an improved method for predicting toxicity of proteins. Brief Bioinform 23: bbac174.

Sherratt, T.N., Roberts, G. & Kassen, R. 2009. Evolutionary stable investment in products that confer both an individual benefit and a public good. Front Biosci 14: 4557–4564.

Simão, F.A., Waterhouse, R.M., Ioannidis, P., Kriventseva, E.V. & Zdobnov, E.M. 2015. BUSCO: assessing genome assembly and annotation completeness with single-copy orthologs. Bioinformatics 31: 3210– 3212.

Smith, W.L., Stern, J.H., Girard, M.G. & Davis, M.P. 2016. Evolution of Venomous Cartilaginous and Ray-Finned Fishes. Integr. Comp. Biol. 56: 950–961.

Smith, W.L. & Wheeler, W.C. 2006. Venom evolution widespread in fishes: a phylogenetic road map for the bioprospecting of piscine venoms. J. Hered. 97: 206–217.

Soneson, C., Love, M.I. & Robinson, M.D. 2015. Differential analyses for RNA-seq: transcript-level estimates improve gene-level inferences. F1000Res. 4: 1521.

Speed, M.P. 1993. Mullerian mimicry and the psychology of predation. Anim. Behav. 45: 571–580. Elsevier BV.

Symula, R., Schulte, R. & Summers, K. 2001. Molecular phylogenetic evidence for a mimetic radiation in Peruvian poison frogs supports a Mullerian mimicry hypothesis. Proc Biol Sci 268: 2415–2421.

Teufel, F., Almagro Armenteros, J.J., Johansen, A.R., Gíslason, M.H., Pihl, S.I., Tsirigos, K.D., et al. 2022. SignalP 6.0 predicts all five types of signal peptides using protein language models. Nat Biotechnol 40: 1023–1025.

von Reumont, B.M., Anderluh, G., Antunes, A., Ayvazyan, N., Beis, D., Caliskan, F., et al. 2022. Modern venomics-Current insights, novel methods, and future perspectives in biological and applied animal venom research. Gigascience 11.

Whitear, M., Zaccone, G., Fasulo, S. & Licata2, A. 1991. Fine structure of the axillary gland in the brown bullhead (Ictalurus nebulosus). J. Zoo 224: 669–676.

Winters, A.E., Chan, W., White, A.M., Van Den Berg, C.P., Garson, M.J. & Cheney, K.L. 2022. Weapons or deterrents? Nudibranch molluscs use distinct ecological modes of chemical defence against predators. J. Anim. Ecol. 91: 831–844.

Winters, A.E., Wilson, N.G., Berg, C.P. van den, How, M.J., Endler, J.A., Marshall, N.J., et al. 2018. Toxicity and taste: unequal chemical defences in a mimicry ring. Proc. Roy. Soc. B, doi: 10.1098/rspb.2018.0457.

Wong, E.S.W. & Belov, K. 2012. Venom evolution through gene duplications. Gene 496: 1–7.

Wright, J.J. 2009. Diversity, phylogenetic distribution, and origins of venomous catfishes. BMC Evol. Biol. 9.

Wright, J.J. 2015. Evolutionary History of Venom Glands in the Siluriformes. In: Evolution of Venomous Animals and Their Toxins, pp. 1–19. Springer, Dordrecht.

Xie, B., Li, X., Lin, Z., Ruan, Z., Wang, M., Liu, J., et al. 2016a. Prediction of Toxin Genes from Chinese Yellow Catfish Based on Transcriptomic and Proteomic Sequencing. Int J Mol Sci 17: 556.

Xie, B., Li, X., Lin, Z., Ruan, Z., Wang, M., Liu, J., et al. 2016b. Prediction of Toxin Genes from Chinese Yellow Catfish Based on Transcriptomic and Proteomic Sequencing. Int. J. Mol. Sci. 17.

Zhu, B., Jin, P., Zhang, Y., Shen, Y., Wang, W. & Li, S. 2023. Genomic and transcriptomic analyses support a silk gland origin of spider venom glands. BMC Biol. 21. BioMed Central.

Ziegman, R., Undheim, E.A.B., Baillie, G., Jones, A. & Alewood, P.F. 2019. Investigation of the estuarine stonefish (Synanceia horrida) venom composition. J. Proteomics 201: 12–26.

