## Supplementary Materials for "Axillary gland gene expression and potency reveal the nature of mimetic relationships in Corydoradinae catfishes"

| **Supplementary Table 1** The BUSCO scores for both the Corydoras and Hoplisoma de novo transcriptome assemblies, indicating quality of assembly. | | | | | | |
| --- | --- | --- | --- | --- | --- | --- |
| Species | Tissue | Complete | Single | Duplicated | Fragmented | Missing |
| *Hoplisoma granti* | Venom gland | 73.1 | 37.6 | 35.5 | 5.4 | 21.5 |
| *Hoplisoma metae* | Venom gland | 81.4 | 26.9 | 54.5 | 3.6 | 15.0 |
| *Hoplisoma tukano* | Venom gland | 67.8 | 41.2 | 26.6 | 5.9 | 26.3 |
| *Corydoras narcissus* | Venom gland | 78.6 | 32.5 | 46.1 | 4.1 | 17.3 |
| *Corydoras desena* | Venom gland | 80.3 | 36.9 | 43.4 | 3.5 | 16.2 |
| *Corydoras simulatus* | Venom gland and Scute tissue | 87.4 | 27.9 | 59.5 | 4.2 | 8.4 |

| **Supplementary Table 2** Post-hoc testing exploring the effect of tissue extract and Corydoradinae genus in of mortality on 24-hour old brine shrimp after a 24-hour incubation period. Asterisks denote significant differences. | | | | | |
| --- | --- | --- | --- | --- | --- |
| **Contrast** | **Tissue / Species** | **Estimate** | **SE** | **z.ratio** | **p.value** |
| **Between Genera** |  |  |  |  |  |
| *Corydoras sp - Hoplisoma sp* | Muscle tissue | 0.213 | 0.804 | 0.265 | 0.791 |
| *Corydoras sp - Hoplisoma sp* | Venom tissue | -0.264 | 0.614 | -0.431 | 0.667 |
| **Within Genus** |  |  |  |  |  |
| Muscle - Venom | *Corydoras sp.* | -2.67 | 0.433 | -6.16 | <.0001*** |
| Muscle - Venom | *Hoplisoma sp.* | -3.15 | 0.49 | -6.424 | <.0001*** |

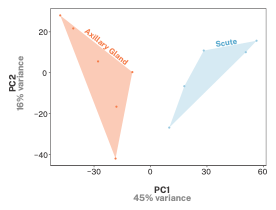

**Supplementary Figure 1** Principal component analysis of RNAseq data from the axillary glands and scute (skin) tissue of Corydoras simulatus.

**Supplementary Figure 2** The toxin network modules represent a group of co-expressed genes thought to be involved in toxin production and regulation, including the majority of identified toxin genes. Downregulation of modules between the axillary gland and scute tissue of C. simulatus. Significant difference is denoted by asterisks (p <0.05, *, p <0.001 **).

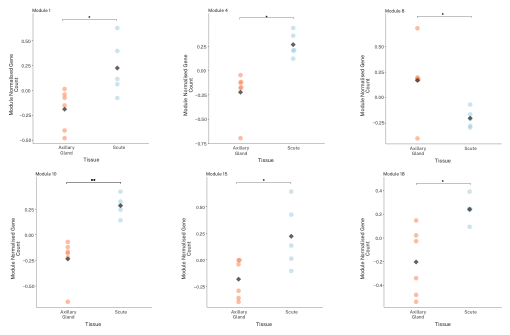

| **Supplementary Table 2** The toxin network module*s* represent a group of co-expressed genes thought to be involved in toxin production and regulation. Negative logfold change denotes upregulation in the axillary gland. | | | | | |
| --- | --- | --- | --- | --- | --- |
|  | Module | Number of Toxin Candidates | Total Gene Number | logFC | adj.P.Val |
|  | ME3 | 216 | 2405 | -0.521 | 0.010 |
|  | ME1 | 74 | 3616 | 0.415 | 0.020 |
|  | ME9 | 70 | 765 | -0.470 | 0.010 |
|  | ME4 | 41 | 1130 | 0.490 | 0.010 |
|  | ME10 | 23 | 578 | 0.519 | 0.010 |
|  | ME8 | 20 | 932 | -0.372 | 0.030 |
|  | ME18 | 12 | 228 | 0.445 | 0.015 |
|  | ME15 | 12 | 273 | 0.405 | 0.020 |

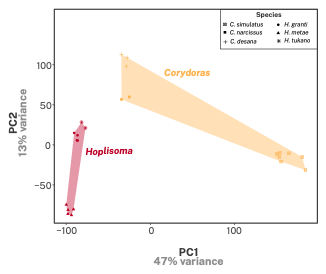

**Supplementary Figure 3** Principal component analysis of RNAseq data from the axillary glands of six species, in two genera. These are H. granti, H. metae, H. tukano from the Hoplisoma and C. narcissus, C. simulatus, C. desana from Corydoras.

| **Supplementary Table 3** The logfold changes associated between orthogroups that are significantly differentially expressed between *Corydoras* and *Hoplisoma* using a phylogenetically controlled framework. | | | |
| --- | --- | --- | --- |
|  | Orthogroup | LogFoldChanges | Genera Upregulated in |
|  | OG0000007 | 0.204 | *Corydoras* |
|  | OG0000028 | 1.757 | *Corydoras* |
|  | OG0000063 | 1.017 | *Corydoras* |
|  | OG0000442 | -3.502 | *Hoplisoma* |
|  | OG0001326 | 3.555 | *Corydoras* |
|  | OG0001490 | 1.154 | *Corydoras* |
|  | OG0002647 | -3.913 | *Hoplisoma* |
|  | OG0003615 | 0.472 | *Corydoras* |
|  | OG0003987 | -1.240 | *Hoplisoma* |
|  | OG0004446 | -1.250 | *Hoplisoma* |
|  | OG0004604 | -0.664 | *Hoplisoma* |
|  | OG0005222 | -1.693 | *Hoplisoma* |
|  | OG0005589 | -3.396 | *Hoplisoma* |
|  | OG0005705 | -1.761 | *Hoplisoma* |
|  | OG0008469 | -2.306 | *Hoplisoma* |
|  | OG0008558 | -2.156 | *Hoplisoma* |
|  | OG0009706 | -2.524 | *Hoplisoma* |
|  | OG0009970 | -0.879 | *Hoplisoma* |
|  | OG0010997 | -2.982 | *Hoplisoma* |
|  | OG0018667 | -1.163 | *Hoplisoma* |
|  | OG0019336 | -4.140 | *Hoplisoma* |
